# A Trade-off Between Developmental Speed and Replication Fidelity Governs the Spontaneous Mutation Rate

**DOI:** 10.1101/2025.10.13.682077

**Authors:** Burak Bulut, Markus Pfenninger

## Abstract

The spontaneous mutation rate (μ) is shaped by two potentially opposing forces: the passage of chronological time, which is inevitably associated with mutation accumulation, and the speed of development, which may compromise replication fidelity. Disentangling these forces has been a major challenge, particularly in ectotherms. Testing the competing predictions of the classic time-dependent Generation Length Hypothesis and the replication-dependent speed-fidelity trade-off model, we experimentally assessed individuals with short (mean 15 days) and long (mean 37.7 days) generation time (GT) in the midge *Chironomus riparius* under constant temperature and estimated the *de novo* mutation rates by whole genome sequencing. We found that long-GT lines accumulated 1.3-fold more mutations per generation (μ/gen), consistent with time-dependent mutagenic processes. Conversely, short-GT lines exhibited a nearly two-fold higher mutation rate per day (μ/day) and a trend towards a transition-biased mutational spectrum (Ts/Tv ratio = 1.28 vs. 0.95), a pattern consistent with a speed-fidelity trade-off in DNA replication. These results suggest that two distinct processes shaped the overall mutation rate. Integrating our data with previous studies and life-history data, we show that the daily mutation rate followed a non-linear relationship with respect to generation time, and that the species’ generation time mode coincides with its minimum. This suggests that the generation time is, amongst other factors, selected to optimise the mutational load by balancing between replication accuracy and developmental speed.

**Teaser Text:** The spontaneous mutation rate (μ) is governed by a fundamental trade-off between the passage of time and the speed of development. We used the midge *Chironomus riparius* to experimentally disentangle these forces. We show that rapid development comes with a cost: compromised DNA replication fidelity and a high daily μ. Conversely, prolonged development accumulates mutations over time. Crucially, the species’ optimal generation time balances these opposing forces, falling precisely at the developmental speed that minimizes the overall mutational load, suggesting that selection for replication accuracy helps shape life-history evolution.

## Introduction

Spontaneous mutations are the ultimate source of all genetic variation, yet the forces that govern their rate (µ) remain a central topic in evolutionary biology (Lynch et al., 2016; Meier et al., 2014). Historically, the mutation rate was often treated as a stable, species-specific constant, a view foundational to the molecular clock hypothesis (Kumar, 2005; Zuckerkandl and Pauling, 1965). This paradigm was later refined by the drift-barrier hypothesis, which proposed that µ is an evolved trait whose lower limit is constrained by the power of selection relative to genetic drift (Lynch et al., 2016), with recent models proposing that this selective pressure is itself dependent on generation time (Zhu et al., 2025). While this framework successfully explains broad-scale variation in µ among species, it is now clear that mutation rates are also highly labile within species, varying between families and in response to factors like parental age (Rahbari et al., 2016), as well as extrinsic environmental pressures (Wu et al., 2024).

There are two competing hypotheses to explain the time-dependent mutation rate variation. The classic Generation Length Hypothesis (GLH) posits that the number of mutations accumulated per generation is primarily a function of chronological time (Lewin and Eyre-Walker, 2025; Wang et al., 2023). This “time-dependent” model suggests that the germline DNA is constantly exposed to damage from endogenous mutagens like reactive oxygen species (ROS), radiation, and spontaneous chemical modification of DNA bases (e.g., deamination, oxidation), with a longer window of exposure leading to a greater mutational load (Salehi et al., 2018; Shokolenko et al., 2009; Srinivas et al., 2018). In contrast, the “replication-dependent” model, often framed as a speed-fidelity trade-off, proposes that the rate of development is a critical factor (Fitzsimmons et al., 2018; Tejada-Arango et al., 2025). Under this model, accelerated cell division and compressed developmental schedules can compromise the accuracy of DNA replication and repair systems, leading to a higher mutation rate per unit of time, not just per cell division or generational passage (Lynch, 2011). These two forces, time-dependent and replication-dependent error, exert distinct pressures on the germline, but experimentally disentangling their relative contributions has remained a major challenge, in part because theoretical models show that both processes can, under different assumptions about repair efficiency, lead to mutation accumulation that scales with cell divisions (Gao et al., 2016), which in turn contributes to a broader scaling conflict where evolutionary predictions differ depending on the time units used (Beichman et al., 2024).

Ectotherms present a particularly acute version of this challenge, as their life-history schedules are inextricably linked to ambient temperature (Carter et al., 2021; Halsey et al., 2015). The recently found U-shaped relationship between temperature and mutation rate in an ectotherm, where µ increased at both cold and warm extremes, exemplifies this complexity (Waldvogel and Pfenninger, 2021). While elevated mutation rates in the heat are readily explained by increased metabolic activity and ROS production, the high mutation rates observed in the cold are puzzling, as they are necessarily associated with prolonged developmental times but not a corresponding increase in metabolic stress (Bulut et al., 2025; Lalouette et al., 2011). This raised the critical question whether the high mutation rate in the cold is a direct consequence of physiological stress, or whether it is an indirect effect of the extended generation time, as predicted by the GLH.

Here, we address these questions disentangling the effects of time and developmental speed on the germline mutation rate and spectrum. We use the non-biting midge, *Chironomus riparius*, an ideal model system that exhibits substantial natural variation in generation time under constant thermal conditions (Foucault et al., 2019). By selecting the earliest and latest reproducing individuals from a single cohort, we created “short” and “long” generation time (GT) lines at a constant, stress-reduced temperature. This design allowed us to isolate the mutational consequences of reproductive timing from those of temperature. This allowed us to simultaneously test the competing predictions of both GLH and speed-fidelity hypothesis, with the former predicting to affect the mutation rate per generation and the mutational spectrum, and the latter the time-dependent mutation rate.

## Material and Methods

### Experimental Design and Rearing

This study employed a modified parent-offspring triad design to assess the effect of reproductive timing on germline mutation rates in *Chironomus riparius*. Because the species’ swarm-mating behaviour prevents direct identification of parents, their genotypes were instead reconstructed from pooled sequencing of their offspring (Oppold and Pfenninger, 2017). The experiment was initiated with 10 egg ropes, ensuring a genetically diverse starting cohort. To control for environmental variation, the resulting larvae were reared at a constant 20°C with a 16:8h light/dark cycle and *ad libitum* feeding (Foucault et al., 2019).

From this mixed cohort, estimated to comprise several hundred individuals, two treatment groups were established based on parental reproductive timing. The “short” GT group was founded from the first ten egg ropes produced by the earliest emerging adults (15 days post-hatching). The “long” GT group was founded from ten egg ropes produced 22-24 days later by late-emerging adults (37-39 days post-hatching) (Sup. Data). Ten replicate lines, descending from ten different parental pairs and thus representing different genetic backgrounds, were established from each group. For each line, the offspring were reared under standardized conditions to the third larval instar stage (L3), at which point ten individual larvae were sampled for individual sequencing, and the remaining siblings (several hundred) were pooled for parental genotype reconstruction. While this pooling approach precludes the identification of individual parents’ genotypes, it is a robust method for accurately reconstructing parental allele occurrences at each locus in species with a large number of offspring, with potential biases mitigated by high sequencing depth and stringent variant filtering criteria.

### Whole-Genome Sequencing and Bioinformatic Analysis

We extracted genomic DNA from the 200 individual L3 larvae (100 per group) and the 20 corresponding sibling pools. Paired-end whole-genome sequencing was conducted on an Illumina platform at Novogene following the methodology of Waldvogel and Pfenninger (2021). Raw read quality was assessed using FastQC. The raw reads then processed using a bioinformatics pipeline based on GATK Best Practices (Van der Auwera et al., 2013). Forward and reverse reads were first merged using PEAR v0.9.6 (Zhang et al., 2014). Merged reads were then mapped to the *C. riparius* reference genome v.4 (Pettrich et al., 2024) using the BWA-MEM algorithm (Li, 2013). To ensure high data quality, duplicate reads were identified and removed using Picard tools v1.123 (“Picard Tools - By Broad Institute,” n.d.), and low-quality reads were filtered with SAMtools (Li et al., 2009).

To identify *de novo* mutations, we analysed the BAM files from each triad (pooled parents and their 10 individual offspring) with the mutation caller accuMUlate (Winter et al., 2017). The resulting candidate mutations were filtered using a custom script with the following criteria: mutation probability ≥ 0.90; probability of exactly one mutation in the triad ≥ 0.90; probability of correct descendant genotype ≥ 0.90; number of mutant reads in the parental pool = 0; mapping quality difference ≤ 2.95; and strand bias or pair-mapping rate difference ≥ 0.05. This multi-layered filtering strategy generates a high-confidence set of *de novo* single-nucleotide mutations (SNMs). Finally, all candidate mutations passing the filtering were manually inspected using the IGV to confirm the mutations (Robinson et al., 2011).

### Statistical Analysis of Mutation Rates and Spectra

We used a Bayesian implementation of a Poisson test (BayesianFirstAid R package) (Bååth, 2014) to estimate the respective median mutation rates along with their 95% high-density interval (HDI) and statistically compare mutation rates between the short and long GT groups. The spontaneous mutation rate (μ) per site per generation was estimated by dividing the number of validated SNMs by the total number of callable genomic sites times the generational passages. The mutation rate per day (μ/day) was calculated by dividing the mutation number by the mean number of callable sites multiplied by the cumulated number of days from egg-laying to reproduction for each line. The mutational spectrum was analysed by calculating the transition-to-transversion (Ts/Tv) ratio and comparing the proportion of transitions between groups with a Bayesian proportion test (Bååth, 2014). To compare the mutation rates obtained from short and long generation times with those from generation times of average length, we combined our data with previously published mutation rate estimates under control conditions for *C. riparius* (Bulut et al., 2024; Rigano et al., 2025). The pooling of these datasets is justified by the highly standardized experimental conditions employed across all studies, including identical rearing temperatures (20°C), light cycles, feeding protocols, and the use of the same bioinformatics pipeline (accuMUlate) with identical parameters for mutation calling. While we cannot completely rule out subtle batch effects from separate experiments, the highly standardized rearing protocols and identical bioinformatic pipelines minimize this possibility and provide the best available data for assessing the mutation rate at intermediate generation times.

### Integration with External Life-History Data

To contextualize our findings in relation to the species’ typical life history, we compiled generation length data from 1,796 control individuals from previous experiments (Bulut et al., 2024; Rigano et al., 2025). This distribution was plotted to visualize the most frequent generation time of *C. riparius* under standard laboratory conditions. As a visual inspection of the plot suggested the mixture of two normal distributions, we used the R-module mixtools (Benaglia et al., 2009) to assess the statistical significance and estimate parameters.

## Results

### Per-Generation Mutation Rate is Higher in the Long-GT Cohort

We successfully established two reproductive cohorts with a mean generation time difference of 22.5 (1.6 s.d.) days. From overall over 1.1 × 10^10^ callable genomic sites, we identified 129 high-confidence *de novo* single-nucleotide mutations (SNMs). Offspring from the short generation group accumulated 57 SNMs, while the long generation group accumulated 72 SNMs. The estimated mutation rate (µ) for the short generation group was 9.40 × 10^−9^ per site per generation (95% HDI = 7.1 × 10^−9^ – 1.2 × 10^−8^). The rate for the long generation group was 1.20 × 10^−8^ per site per generation (95% HDI = 9.7 × 10^−9^ – 1.5 × 10^−8^). This represents, with a posterior probability of 93.4% an increase (median = 1.3-fold increase) in the germline mutation rate in the offspring of parents that reproduced later (Figure 1A).

**Figure 1.**
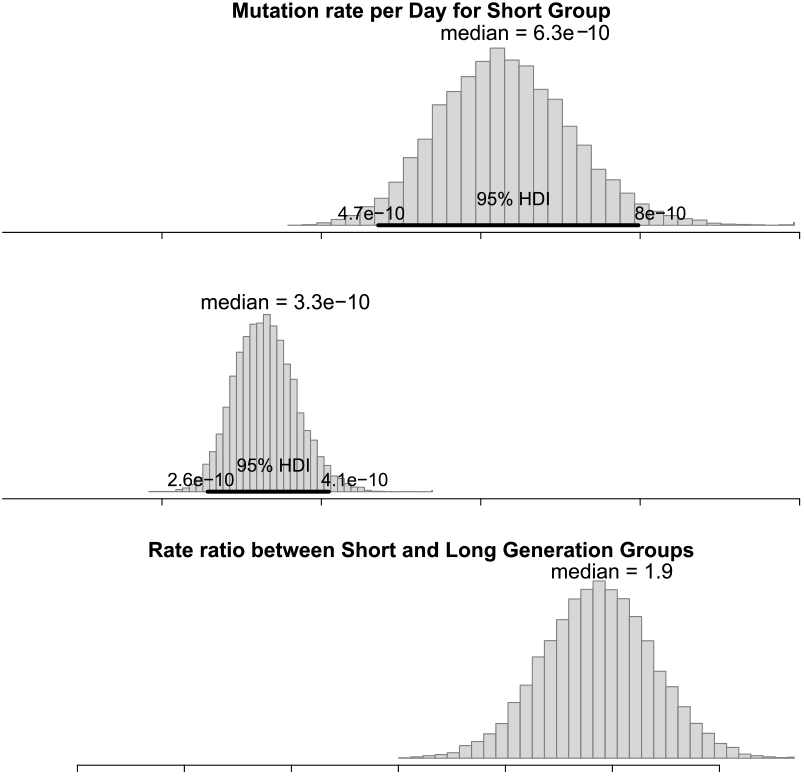
The effect of generation time on the per-generation mutation rate. (A) Posterior probability distribution of the rate ratio (λ_long / λ_short) of the per-generation mutation rate between the long and short generation time cohorts. The median rate ratio is 1.3, indicating a 1.3-fold higher rate in the long-GT group. The 95% Highest Density Interval (HDI) is [0.94, 1.9]. The posterior probability that the rate in the long-GT group is higher than in the short-GT group (ratio > 1) is 94.2%. (B) In contrast, the per-day mutation rate ratio between short and long generation groups has a median of 1.9, signifying a 1.9-fold higher rate in the short group. The 95% HDI is [1.4, 2.8]. The posterior probability that the rate in the short group is higher than in the long group (ratio > 1) is nearly 100%.

### Per-Day Mutation Rate is Higher in the Short-GT Cohort

In contrast to the per-generation rate, the mutation rate per day was nearly two-fold higher in the short-GT cohort. The estimated daily rate (μ/day) for the short-GT group was 6.2 × 10^−10^ per site per day (95% HDI = 4.7 × 10^−10^ – 7.9 × 10^−10^), while the rate for the long-GT group was 3.3 × 10^−10^ per site per day (95% HDI = 2.5 × 10^−10^ – 4 × 10^−10^). Bayesian analysis showed a posterior probability of 99.8% that the daily rate in the short-GT group was higher than in the long-GT group, with a median rate ratio of 1.9 (Figure 1B).

### Mutational Spectra Differ Between Generation Time Cohorts

There was evidence that the mutational spectrum was shifted between the two cohorts. The short-generation group exhibited a transition-to-transversion (Ts/Tv) ratio of 1.28, while the long-generation cohort had a Ts/Tv ratio of 0.95 (Figure 2A). A Bayesian proportion test indicated an 84.1% posterior probability that the proportion of transitions was higher in the short-generation group (estimated group difference 95% CI: -0.081 to 0.25). This shift was primarily driven by the relative frequencies of specific mutation types. A↔G transitions were significantly more frequent in the short-generation group (95% CI for difference: -0.009 to 0.31; posterior probability = 0.965). Conversely, C↔T transitions showed a trend towards being more frequent in the long-generation group (95% CI for difference: -0.25 to 0.081; posterior probability = 0.882) (Figure 2B).

**Figure 2.**
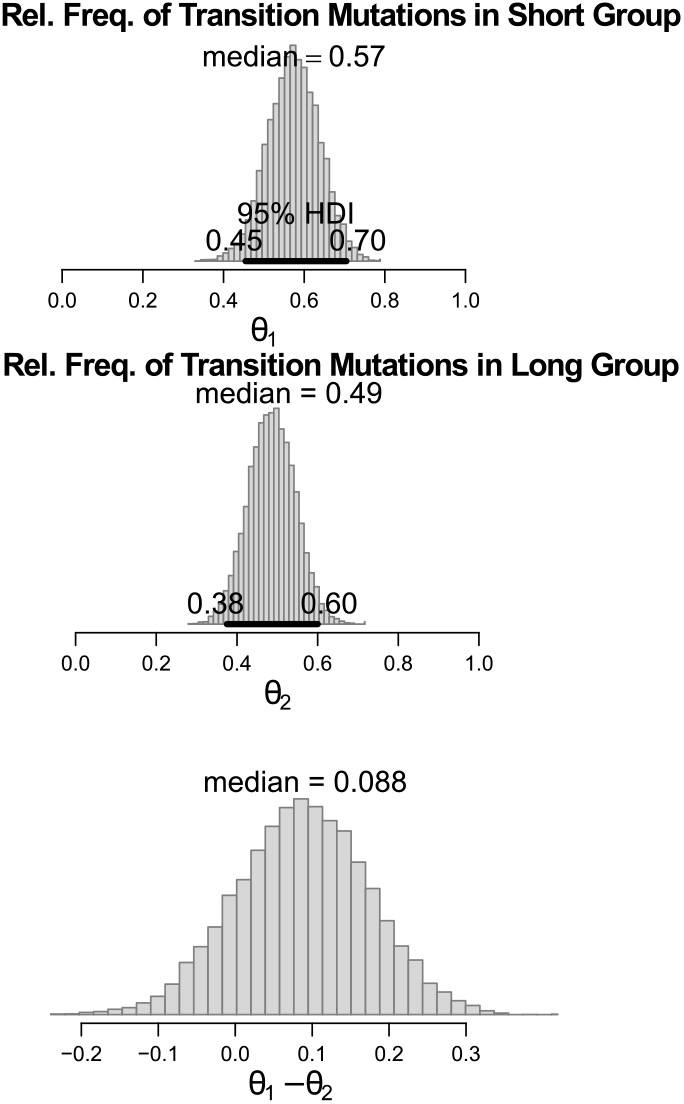
Mutational spectrum of the short and long generation time cohorts. (A) Heatmap showing the absolute counts of the six single-nucleotide mutation types. The y-axis categorizes mutation types, with transitions (A↔G, C↔T) in the top two rows and transversions below. The x-axis separates the two experimental groups. The spectrum is biased towards transitions in the short-GT group (Ts/Tv = 1.28) and is more balanced in the long-GT group (Ts/Tv = 0.95), with A↔G transitions being significantly more frequent in the short-GT cohort. (B) The relative frequencies of specific mutation types. Bayesian proportion test indicated an 84.1% posterior probability that the proportion of transition mutations was higher in the short group (95% CI for difference: -0.081, 0.25).

### The Relationship Between Daily µ and Generation Time is Non-Linear

Using the mutation rates inferred under stress-free control conditions for intermediate generation length in other studies (Bulut et al., 2024; Rigano et al., 2025) and unpublished data suggested that the relationship between generation time and mutation rate is non-linear, as evidenced by the non-overlap of the 95% HDI intervals to both short and long GT groups (Figure 3). To visualize the potential non-linear relationship between mutation rate and generation time, we plotted a quadratic curve, which suggested a nadir around 26 days for the per day rate and 29 days per generation (Figure 3A + B).

**Figure 3.**
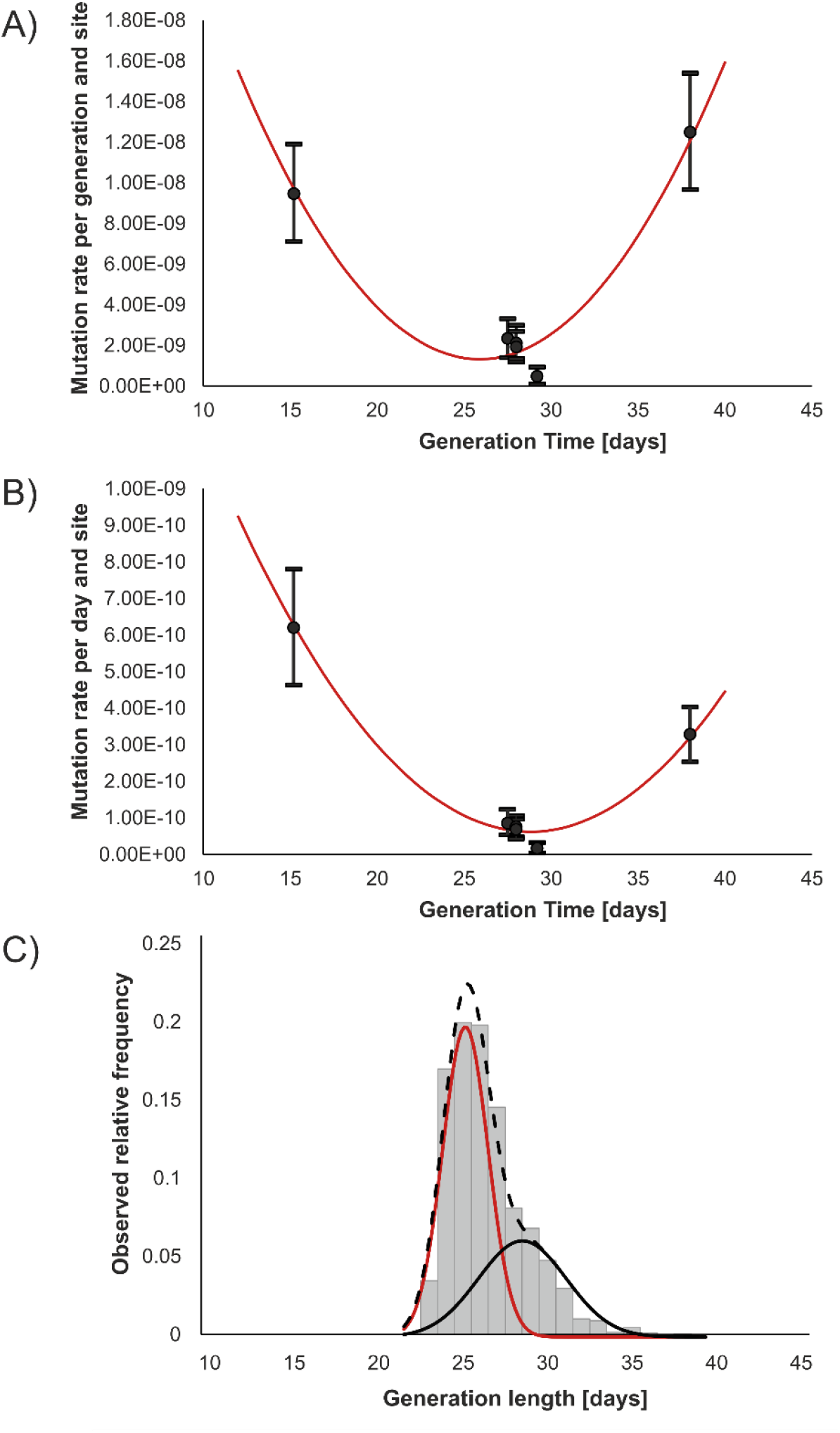
Mutation rate as a function of generation length. A) Mutation rate per generation. B) Mutations per day. Error bars correspond to 95% HDI. The solid red line shows the best-fit quadratic functions. C) Frequency distribution of 1796 generation time measurements. The red and black solid lines represent the inferred distinct underlying normal distributions, and the dashed line is the inferred mixed distribution.

### Modal Population Generation Time Aligns with the Minimum Daily µ

An analysis of 1,796 individuals from control conditions of previous experiments revealed that the mean generation time for *C. riparius* under standard laboratory conditions was 26.4 (s.d. 2.3) days (Figure 3C). This peak in reproductive timing fell between the minima for both per-day and per-generation mutation rates. However, the generation time data were much better described by a mixture of two normal distributions (p single mode < 1 x 10^-16^). The estimated parameters of these two distributions were a mean generation time of 25.3 +/-1.2 days for the first mode and 28.3 +/-2.3 days for the second, with relative proportions of 0.62 and 0.38, respectively.

## Discussion

Our study provides direct experimental evidence that the germline mutation rate is shaped by a dynamic balance between two distinct, independent forces tied to generation time. By isolating generation time from temperature in *Chironomus riparius*, we demonstrate that a “rate-dependent” mutational regime, driven by a speed-fidelity trade-off, dominates at short generation times, while a “time-dependent” regime, governed by mutation accumulation, prevails at long generation times. This finding helps resolve the simple dichotomy between time and replication-dependent models of mutagenesis. It provides empirical support for theoretical frameworks predicting that mutation accumulation depends on the interplay between mutation rates and repair efficiency (Gao et al., 2016). While the current study is restricted to our model system, there is little reason to believe that these processes should act principally differently in other ectotherms.

### A Speed-fidelity Trade-off Drives Mutagenesis at Short Generation Times

The mutational signature of the short-GT cohort is a pattern consistent with a speed-fidelity trade-off. These individuals exhibited a nearly two-fold higher mutation rate per day than the long-GT cohort, likely a direct cost of their compressed developmental schedule. This interpretation is strongly supported by the observed shift in the mutational spectrum; the increased observed transition bias (Ts/Tv ratio = 1.28) is a molecular signature frequently associated with replicative polymerase errors, which are expected to occur more often when developmental processes are accelerated (Fitzsimmons et al., 2018; Kunkel, 2004; Lynch, 2010). This happens potentially by altering DNA replication dynamics; for instance, mutation rates are known to be higher in genomic regions that replicate late in S-phase (the DNA synthesis phase of the cell cycle), a state that may be more prevalent under temporal pressure (Stamatoyannopoulos et al., 2009). In systems ranging from viruses to yeast, such trade-offs are a known constraint on rapid growth (Liu and Zhang, 2021; Regoes et al., 2013), and our results demonstrate this principle operating in the germline of a complex animal, providing strong evidence for a rate-dependent mutational load due to precocious reproduction. While the trend towards a higher transition bias in the short-GT cohort did not reach the highest levels of statistical certainty, it is consistent with the molecular signatures of replicative errors and supports our mechanistic interpretation. A larger sample of mutations would be needed to resolve the finer details of the respective mutational spectra.

### Time-dependent DNA Decay Increases the Mutational Load at Long Generation Times

Conversely, the long-GT cohort exemplifies the classic “Generation Length Hypothesis”, but with a critical temporal nuance. These lines accumulated a 1.3-fold higher mutation rate per generation, confirming that a longer chronological window allows for a greater total accumulation of mutations (Latta et al., 2013; Thomas et al., 2010). Despite this higher generational total, however, the daily influx of mutations was significantly lower than in the short-GT group. The mutational spectrum gave a hint on the resolution of this apparent paradox. The shift to a lower Ts/Tv ratio (0.95) is indicative of a greater relative contribution from time-dependent DNA damage, such as oxidative lesions that are known to produce transversions (La et al., 2023; Wang and Obbard, 2023). This indicates that while the chronological clock of mutation accumulation is always ticking, its daily rate is substantially lower than the rate of error introduction during rapid replication. This demonstrates that while the chronological ‘clock’ is a persistent source of mutations, its per-day impact on the total mutation rate is small compared to the errors introduced by the ‘replication fidelity clock’ when development is accelerated.

### The Population’s Mean Generation Time Coincides with the Mutational Minimum

Bayesian analysis showed that the 95% HDI for the mutation rates at intermediate generation lengths did not overlap with those from either the short or the long generation time groups. This indicated a posterior probability of nearly 100% that the mutation rate at the nadir is truly lower than at the extremes. These distinct mutational regimes predict that the daily mutational burden should be a non-linear (perhaps quadratic) function of generation time, but the evolutionary relevance of this prediction hinges on where the species’ natural life history falls on this landscape.

Our analysis of generation time over 1,700 individuals from earlier studies (see above) revealed that the mean of the generation time distribution for *C. riparius* under these experimental conditions lies between the nadirs of both the daily mutation rate (Figure 3B) and the generational mutation rate (Figure 3A). Even more interestingly, the analysis of the generation time distribution revealed that it is significantly better described by a mixture of two normal distributions (Figure 3C). The two-modal distribution is consistent with a model where an allele of large effect segregates in the population in addition to polygenic variation (Jain and Stephan, 2015). The polygenic basis of the trait generation time implies that there is phenotypic variation of this trait in natural populations (Barghi et al., 2019) if not all contributing loci are fixed. The non-linear reaction norm of the mutation rate in relation to generation time found here therefore means that the mean mutation rate in natural populations is likely to be higher than that measured in the laboratory at optimal generation time, regardless of other natural (e.g. Waldvogel and Pfenninger, 2021) and anthropogenic (e.g. Bulut et al., 2024; Rigano et al., 2025) factors that may increase it.

In principle, earlier reproduction should increase individual fitness by producing more direct and later generation offspring faster, particularly in a species with a predominantly semelparous life history like *C. riparius*, where fitness is largely determined by a single, successful reproductive event following maturation (Brommer et al., 2002). If there is genetic variation in developmental speed, there should be a selective pressure towards early reproduction. Experimental evolution on developmental time clearly showed a strong selective response, indicating a broad, polygenic basis for generation time variation (Caliendo 2025). However, a shorter developmental time is inevitably associated with costs, and the mutational load associated with a higher mutation rate is likely one of them (Leslie et al., 2017). While some theories propose that higher mutation rates could be adaptive under stressful conditions by increasing “evolvability” (Metzgar and Wills, 2000), theoretical and empirical evidence show that mutational load tends to decrease fitness (Grossen and Ramakrishnan, 2024; Lynch et al., 2016). Our data is in line with the latter view: the population mean of generation time coincides with the minima of both daily and generational mutation rate. This is consistent with the standard view that selection acts powerfully to reduce the influx of predominantly deleterious mutations as far as drift allows (Baer et al., 2007; Lynch et al., 2016). At this evolutionarily favoured optimum, development is slow enough to maintain high replication fidelity, mitigating the costs of speed, yet rapid enough to avoid the substantial accumulation of time-dependent mutations, mitigating the costs of time. A first step towards demonstrating this would be to prove that individuals with extreme generation times are less fit than individuals at the supposed optimum. It would also be interesting to see whether individual mutation rates along the generation time gradient exhibit dynamic cryptic genetic variation, as is the case with temperature (Pfenninger et al., 2025). As a side effect, our findings also suggest that the increase in mutation rate in the cold (Waldvogel and Pfenninger, 2021) is primarily an indirect effect of extended generation time rather than a direct consequence of low-temperature stress, a conclusion consistent with the finding of only a minimal increase in ROS under this temperature (Bulut et al., 2025).

### Implications for Life-history Evolution

The current view of mutation rate evolution acknowledges that while ultimate limits may be set by the drift-barrier (Lynch et al., 2016), the proximate rate is a highly labile trait shaped by an organism’s internal and external environment (Sharp and Agrawal, 2018). Our dual-mechanism optimization model provides a novel hypothesis for an environmentally independent part of this lability by suggesting a direct link between an organism’s generation time and its mutational load.

## Acknowledgements

We gratefully acknowledge the support of this study by a grant of the Deutsche Forschungsgemeinschaft (PF 390/15-1) to MP.

## Data Availability Statement

All nucleotide sequences used in this study are deposited in the European Nucleotide Archive (ENA) under the project number PRJEB89193. Raw data for the analyses were made available at Zenodo doi: **doi.org/10.5281/zenodo.17241504**

